# Generation of Xylose-inducible promoter tools for *Pseudomonas* species and their use in implicating a role for the Type II secretion system protein XcpQ in inhibition of corneal epithelial wound closure

**DOI:** 10.1101/2020.01.31.929794

**Authors:** Jake D. Callaghan, Nicholas A. Stella, Kara M. Lehner, Benjamin R. Treat, Kimberly M. Brothers, Anthony J. St. Leger, Robert M. Q. Shanks

## Abstract

Tunable control of gene expression is an invaluable tool for biological experiments. In this study, we describe a new xylose-inducible promoter system and evaluate it in both *Pseudomonas aeruginosa* and *P. fluorescens*. The *P*_*xut*_ promoter derived from the *P. flurorescens xut* operon was incorporated into a broad host-range pBBR1-based plasmid and compared to the *Escherichia coli*-derived *P*_*BAD*_ promoter using *gfp* as a reporter. GFP-fluorescence from the *P*_*xut*_ promoter was inducible in both *Pseudomonas* species, but not in *E. coli,* which may facilitate cloning of toxic genes using *E. coli* to generate plasmids. The *P*_*xut*_ promoter was expressed at a lower inducer concentration than *P*_*BAD*_ in *P. fluorescens* and higher *gfp* levels were achieved using *P*_*xut*_. Flow cytometry analysis indicated that *P*_*xut*_ was more leaky than *P*_*BAD*_ in the tested *Pseudomonas* species, but was expressed in a higher proportion of cells when induced. D-xylose did not support growth of *P. aeruginosa* or *P. fluorescens* as a sole carbon source and is less expensive than many other commonly used inducers which could facilitate large scale applications. The efficacy of this system aided in demonstrating a role for the *P. aeruginosa* type II secretion system gene from *xcpQ* in bacterial inhibition of corneal epithelial cell wound closure. This study introduces a new inducible promoter system for gene expression for use in *Pseudomonas* species.

**Importance:** *Pseudomonas* species are enormously important in human infections, biotechnology, and as a model system for interrogating basic science questions. In this study we have developed a xylose-inducible promoter system and evaluated it in *P. aeruginosa* and *P. fluorescens* and found it to be suitable for the strong induction of gene expression. Furthermore, we have demonstrated its efficacy in controlled gene expression to show that a type 2 secretion system protein from *P. aeruginosa*, XcpQ, is important for host-pathogen interactions in a corneal wound closure model.

## INTRODUCTION

Species of the bacterial genus *Pseudomonas* are of exceptional importance, not only as an infectious agent for a broad range of organisms including humans and plants (1–4), but also for the valuable role species of this genus play in biotechnology and basic science (5, 6). Important tools in both biotechnology and basic science are inducible gene expression systems that enable induced expression or repressed expression in the presence of an effector molecule.

Inducible plasmid systems for pseudomonads include the L-arabinose inducible, AraC regulated *P*_*BAD*_ promoter from *Escherichia coli*, which is inducible in *P. aeruginosa* (7–9). Whereas this promoter has proven useful for gene expression in numerous studies, it has been demonstrated to be leaky without induction in *P. aeruginosa* (10). Other plasmid-based promoter systems inducible in *P. aeruginosa* systems include *lacI^q^*-*Ptac* and the *rhaSR*-*P_rhaB_* promoter (10, 11).

The goal of this study was to develop a new inducible promoter system for *Pseudomonas* species using pseudomonad-derived DNA. This study describes the evaluation of the *P. fluorescens xutR*-*P_xutA_* promoter system, here noted as *P*_*xut*_, in both *P. aeruginosa* and *P. fluorescens*. In addition, we use this system to interrogate the ability of *P. aeruginosa* host-pathogen interactions, particularly, the mechanisms underlying bacterial inhibition of corneal epithelial cell wound closure (12).

## MATERIALS and METHODS

### Strains, media, and growth conditions

Bacterial strains and plasmids were listed in Table 1. Bacteria were grown in lysogeny broth (LB) (13), and aerated on a TC-7 tissue culture roller (New Brunswick, Inc). *P. aeruginosa* strains are listed in Table 1. Gentamicin was used at 10 μg/ml in *E. coli* and 30μg/ml in *Pseudomonas* species to select for plasmids. Bacterial culture density was measured with a 1 cm cuvette using a SpectraMax M3 plate reader, except where noted below.

**Table 1.** Strains and plasmids used in this study.

| Strain or plasmid | Description | Reference or Source |
| --- | --- | --- |
| S17-1 λ-pir | *E. coli* laboratory strain | (37) |
| PA14 | wild-type *P. aeruginosa* UCBPP-PA14 | (38) |
| Pf0-1 | *P. fluorescens* wild-type strain | (39) |
| pMQ80 | source of *gfpmut3* | (8) |
| pMQ132 | Shuttle vector, pBBR1, *P<sub>lac</sub>-lacZ*, *aacC-1* | (11) |
| pMQ578 | pMQ132 with *xutR- P<sub>xut</sub>-gfp* replacing *P<sub>lac</sub>-lacZ* | This study |
| pMQ630 | pMQ132 with *P<sub>BAD</sub>-gfp* replacing *P<sub>lac</sub>-lacZ* | This study |
| pMQ643 | pMQ578 with *tdtomato* replacing *gfp* | This study |
| pMQ644 | pMQ578 with *xcpQ* from PA14 replacing *gfp* | This study |
| pMQ650 | pMQ578 with multicloning site replacing *gfp* | This study |
| pMQ652 | pMQ650 with yeast replicon removed | This study |

### Plasmid construction

Plasmids are listed in Table 1 and oligonucleotide primers and synthetic DNA sequences are listed in Table 2. Plasmids were made using yeast homologous recombination (8) except pMQ652 and verified by PCR and sequencing of junctions. The *xutR* gene and intergenic region between *xutR* and *xutA* from the *Pseudomonas fluorescens* Pf0-1 genome was cloned along with *gfpmut3* (*gfp*) (14) from pMQ80 (8) to replace *P_lac_* and *lacZα* from pMQ132 (11). The resulting plasmid, pMQ578, and all plasmids made in this study are listed in Table 1. One variant of pMQ578 was made in which *gfp* was replaced with *tdtomato* from pMQ414 (15) using primers 4127 and 4218 and named pMQ643. For another, pMQ650, an artificial DNA sequence containing six restriction enzyme sites (EcoRI, SalI, BamH1, SpeI, SphI, and HindIII) and a sequence with stop codons in three frames was introduced using a 499 bp long double stranded DNA sequence listed as primer number 4068 (gBlock, IDT, Inc). Lastly, pMQ652 was made by digestion of pMQ650 with SspI and StuI, which excises a 1.8 kba region containing the yeast replicon and *URA3* gene, followed by recircularization with T4 DNA ligase.

**Table 2.**
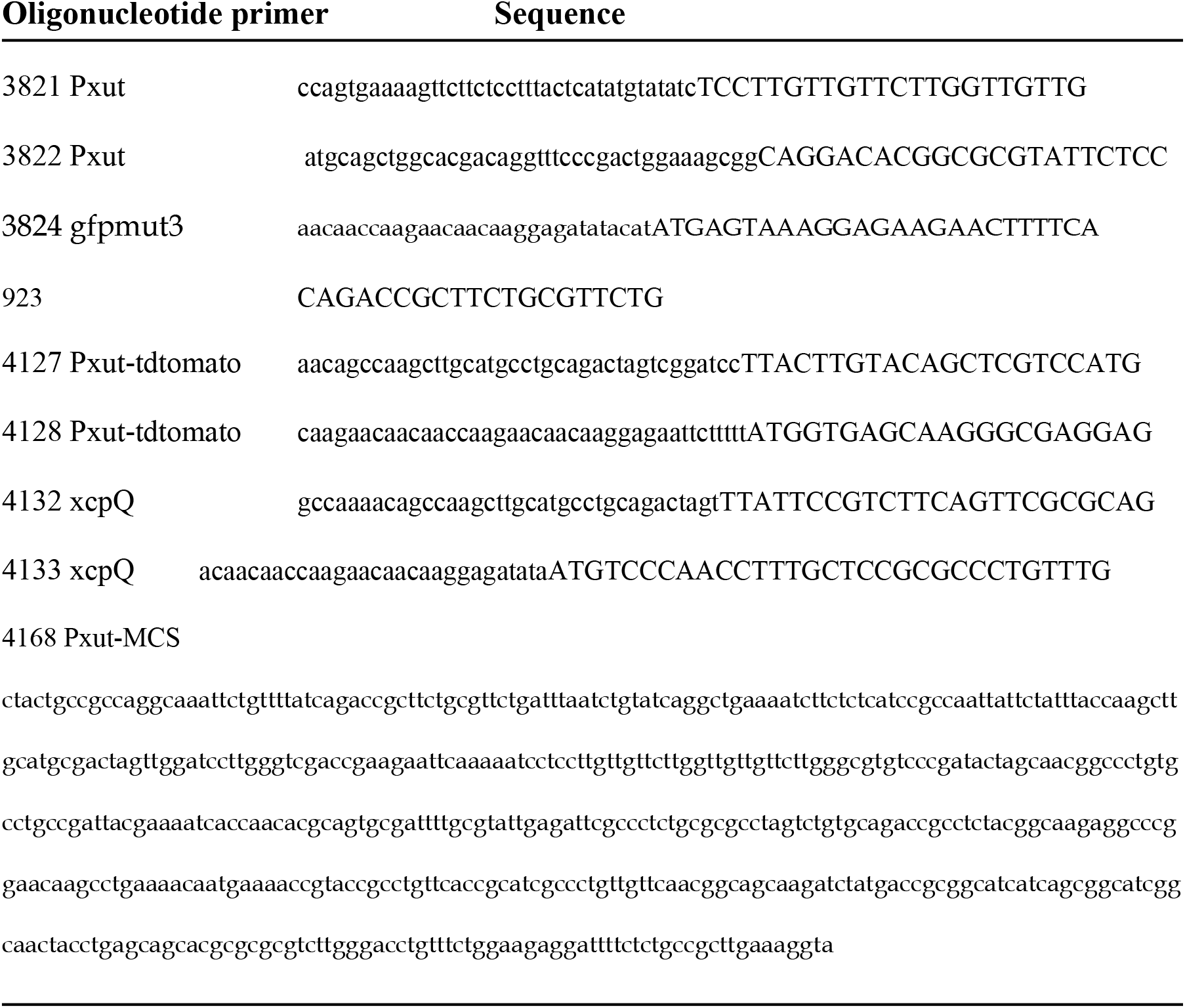
Primers and synthetic DNA used in this study.

### Fluorescence assays

Bacteria were grown as above and at indicated time points samples were obtained for analysis. Aliquots (150 μl) were read in 96 well plates with black opaque sides and clear bottoms (Thermo Scientific Nunc 165305). GFP and tdtomato fluorescence and optical density at 600 nm was read with a BioTek Synergy 2 plate reader using 485/20 excitation and 516/20 emission filters for GFP and 545/40 excitation and 590/20 emission and filters, respectively.

### Flow cytometry analysis of GFP fluorescence

Bacteria were grown overnight with 10 mM inducer, adjusted to OD_600_ = 2,washed twice, and adjusted to OD_600_=0.02 in phosphate buffered saline (PBS) that had been filtered with a 0.22 μm filter Flow cytometric analysis was performed on a CytoFLEX LX instrument (Beckman Coulter, USA). Filtered PBS was used as negative control for gating bacteria by forward scatter (FSC) and side scatter (SSC). Approximately, 10^5^ cells were analyzed per genotype, and the experiments were repeated on five occasions. Bacteria of each species without the fluorescent plasmid were used as negative control to determine FSC threshold, and additionally for background fluorescence cutoff. Software analysis was performed using FlowJo™ version 10 (Becton Dickinson, USA).

### Corneal cell migration analysis

In vitro epithelial cell wound closure assays were performed as previously described (12) using human corneal limbal epithelial (HCLE) cells (16) and a commercial wound healing assay kit (ORIS™ Platypus Technologies, LLC # CMAU101). After stratified cell layers of HCLE cells were established (17), they were challenged with 50 μl of normalized bacterial culture filtrates or LB medium that were added to 100 μl of culture media overlaying the cells. Tissue culture medium consisted of keratinocyte serum-free medium (KSFM) (Gibco cataolog number 10724-011) supplemented with bovine pituitary extract (25 μg/ml), epidermal growth factor (0.2 ng/ml), penicillin and streptomycin at 100 μg/ml. Bacterial culture filtrates were made using overnight cultures of *P. aeruginosa* grown in LB medium with or without xylose and antibiotic and was incubated at 30°C. Cultures were adjusted to OD_600_=1.0 using fresh LB medium and filtered with a 0.22 μm filter (Mellex-GV PVDF). After 20-24 hours HCLE cells were rinsed with PBS and stained using 0.5 μM Calcein AM (Invitrogen catalog number C3099) for 15 minutes. Cells were imaged using an Olympus Fluoview FV-1000 laser scanner confocal microscope with a 4X (0.3) NA objective and analyzed with Fluoview image viewing software.

### Bacterial length analysis

Bacteria were grown in LB medium with and without inducer (arabinose or xylose at 10 mM) for 24 hours. Bacteria were then imaged as noted above, but using a 60X objective. Bacteria were measured using ImageJ software and at least 50 cells were counted from two different experiments per group.

### Statistical analysis

Experiments were performed at least three times and data were analyzed with GraphPad Prism software using ANOVA, two-way ANOVA, and Student’s T-tests.

## RESULTS

### Identification and cloning of a putative xylose-inducible promoter from *P. fluorescens*

A putative xylose metabolism transcription factor gene, Pf101_2304 was identified in the *P. fluorescens* strain Pf0-1 genome by BLAST analysis (18). Recently, the homolog of this gene in *P. fluorescens* strain SBW25 gene was demonstrated to be an activator of a xylose metabolism operon and was named XutR by Liu, et al (19). The XutR protein was demonstrated to directly and positively regulate transcriptional expression of the adjacent gene, *xutA* (Figure 1A), in a xylose-dependent manner, and bound to a conserved operator site (AAAATC-N15-GATTTTT) upstream of *xutA* (19) (Figure 1B). In the SBW25 genome, the intergenic region between *xutR* and *xutA* is 139 bp in length, whereas it is 182-188 in *P. fluorescens* strain Pf0-1 depending upon whether a TTG or ATG for *xutA* is the start codon (Figure 1B). Other differences in the intergenic region between strain SBW25 and Pf0-1 include a direct repeat of CCAAGAACAACAA just upstream of the ribosome binding site in Pf0-1 that is present in a single copy in SBW25. The *xutR* and intergenic region with the promoter for *xutA*, noted here as *P*_*xut*_, were cloned in a broad host range pBBR1-based plasmid, pMQ132 (11) with *gfp* placed under control of *P*_*xut*_ (Table 1). DNA sequence of the *xutR* and *P*_*xut*_ region from pMQ578 was deposited in GenBank (accession number MN857504).

**Figure 1.**
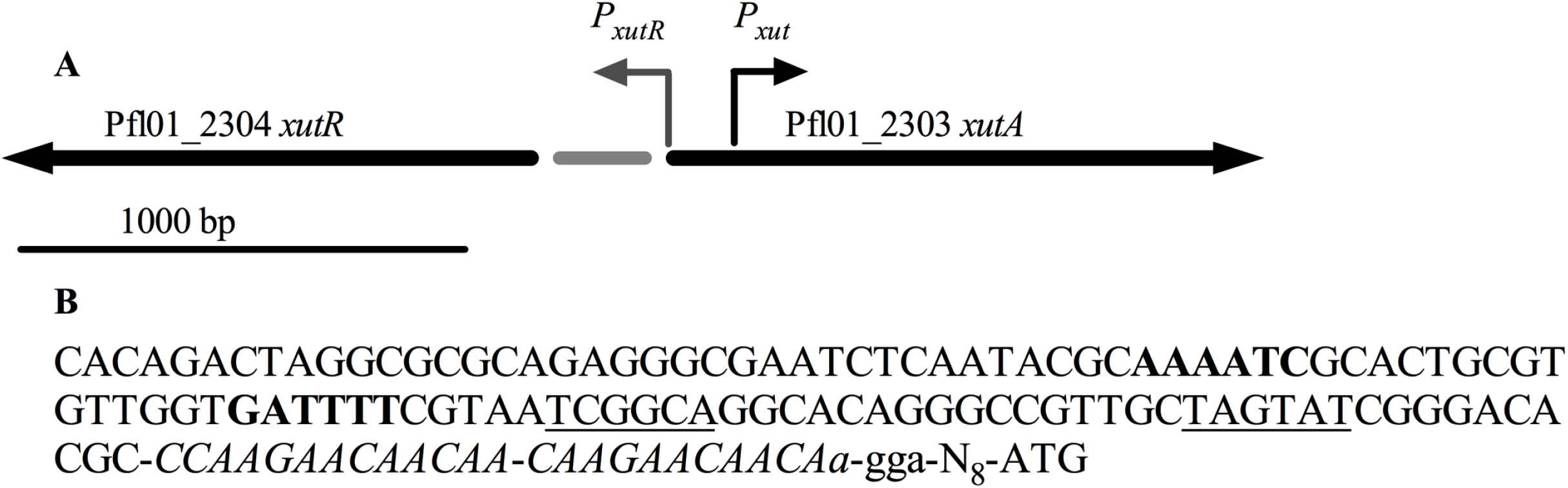
Xylose repressor genomic region from *P. fluorescens* strain Pf0l. **A.** Genetic map of the *xutR-xutA* region. The gray bar indicates the 188 bp intergenic region. **B.** DNA sequence of the *P*_*xut*_ promoter upstream of Pfl01_2303. The conserved operator inverted repeats are shown in bold and the −35 and −10 regions are for the *xutA* promoter (*P*_*xut*_) are underlined. A direct repeat of 13 base pairs is shown in italic letters. A putitive ribosome binding site 9-12 base pairs upstream of the start codon is in lower case.

The pMQ578 plasmid has an RP4 *oriT* sequence for conjugal transfer, the *aacC-1* gentamicin resistance marker, and a selectable marker and replicon for *Saccharomyces cerevisiae* to allow for yeast in vivo cloning (8). We also replaced the *P*_*xut*_ promoter in pMQ578 with the *E. coli P_BAD_* promoter to generate pMQ630; this plasmid was used as a comparison promoter, since the *P*_*BAD*_ promoter is frequently used for gene expression in *E. coli* and *Pseudomonas* species. The resulting plasmids were used to characterize *P*_*xut*_ expression in *E. coli* and two species of *Pseudomonas*.

### Characterization of *P*_*xut*_ expression in *E. coli*, *P. aeruginosa*, and *P. fluorescens*

To evaluate *P*_*xut*_ and *P*_*BAD*_ expression, pMQ578 and pMQ630 were introduced into *E. coli* strain S17-1, *P. aeruginosa* strain PA14, and *P. fluorescens* strain Pf0-1. Increasing doses of L-arabinose and D-xylose were added to cultures (OD_600_ = 0.01), and cultures were analyzed for GFP fluorescence using a fluorometer after 25 hours of incubation at 30°C. After induction, with *E. coli*, there was no fluorescence measured with *P_xut_-gfp*, whereas, *P*_*BAD*_ was inducible at low concentrations (0.1 mM) of L-arabinose (Figure 2A). In *P. aeruginosa*, both *P*_*xut*_ and *P*_*BAD*_-based expression was dose dependent with the *P*_*xut*_ promoter being slightly stronger at intermediate inducer doses (Figure 2B). With *P. fluorescens*, both promoters were inducible, but the *P*_*xut*_ promoter was inducible at lower concentrations and was stronger at all doses (Figure 2C).

**Figure 2.**
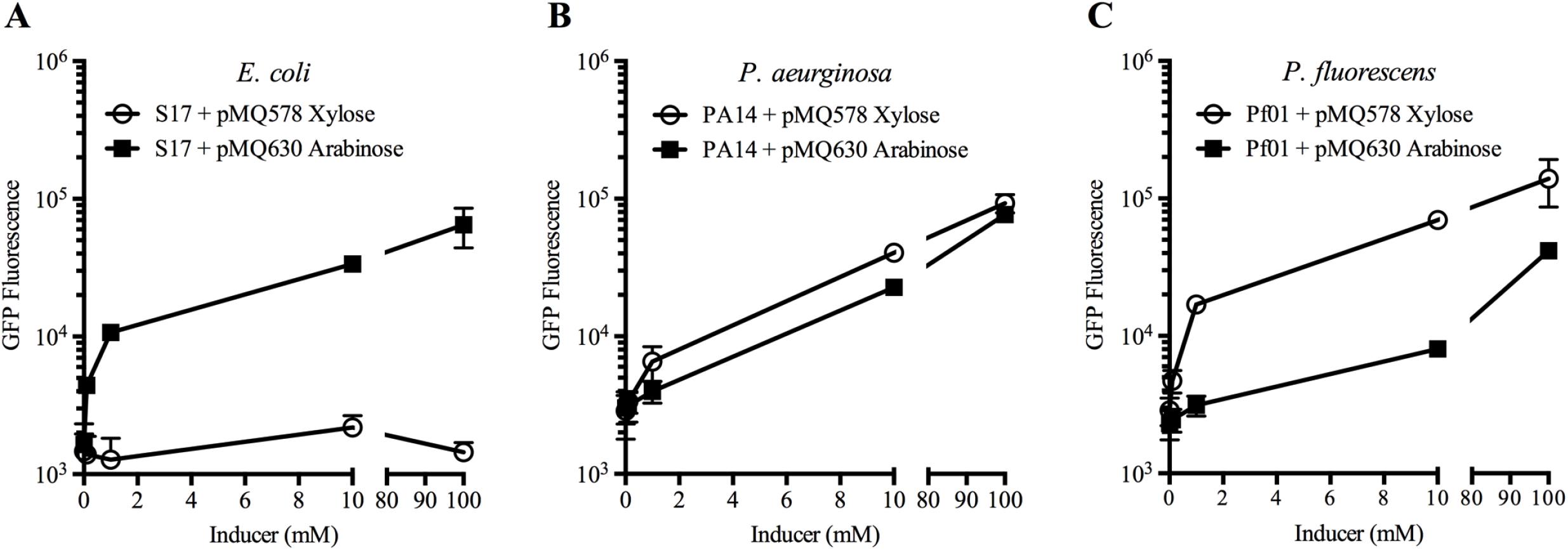
Comparison of *P*_*BAD*_ and *P*_*xut*_ promoter in *E. coli* and *Pseudomonas* species. Expression of *gfpmut3* from the two different promoters in a pBBRl-based plasmid. Cultures were prepared in LB medium with a range of inducer concentrations and grown at 30°C for 25 hours. Mean and SD are given, n≥5 independent cultures. *P*_*xut*_ was not inducible in *E. coli* and was inducible at lower inducer conditions in *P. fluorescens.* The pMQ578 plasmid has *P*_*xut*_-*gfp*, and pMQ630 has *P*_*BAD*_-*gfp*.

Expression of both promoters was tested over time using 10 mM inducer (Figure 3A-C). Cultures with inducer were started at OD_600_ = 0.01 and allowed to grow at 30°C. Samples were removed and analyzed for GFP fluorescence over time. GFP fluorescence was largely undetected until 6 hours post induction, and was maximal at the final time point (25 hours). The highest fluorescence levels were observed with Pf0-1 with *P*_*xut*_ (pMQ578) at 25 hours (Figure 3C).

**Figure 3.**
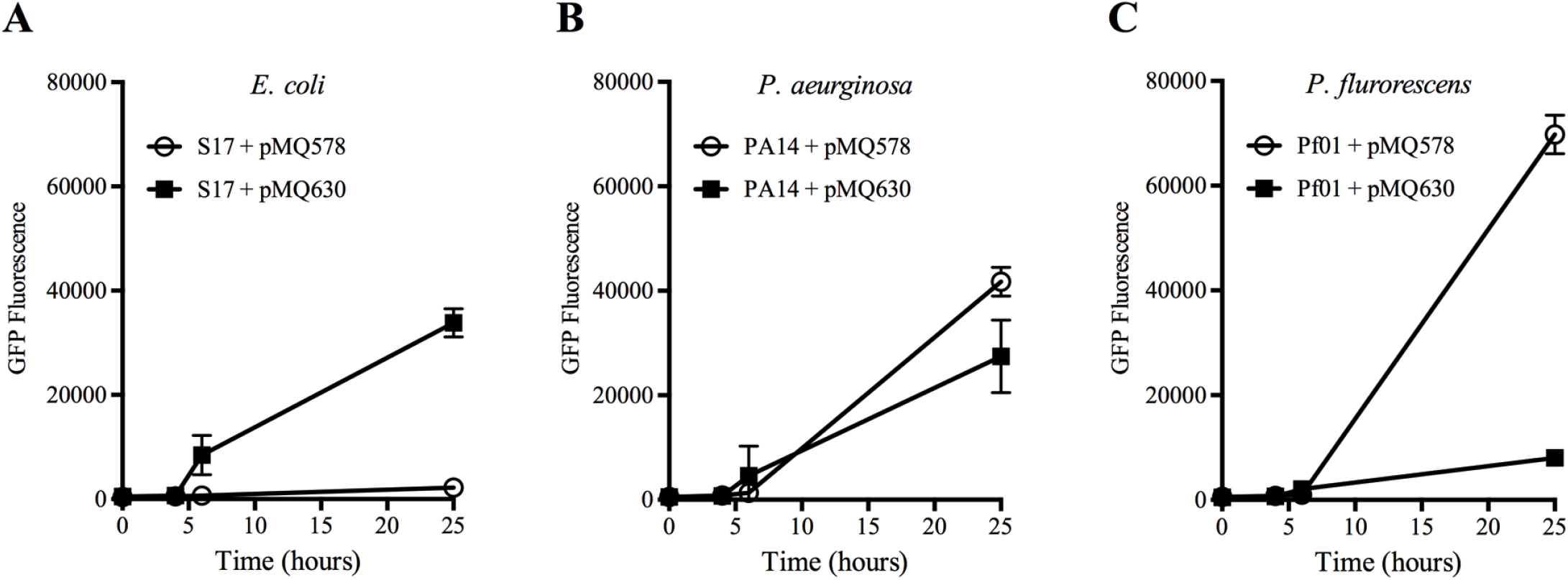
Expression of *P*_*BAD*_ and *P*_*xut*_ promoter in *Pseudomonas* species over time with 10 mM inducer. Expression of *gfpmut3* over time from the two different promoters in a pBBRl-based plasmid. Cultures were prepared in LB medium with inducer (10 mM) and grown at 30°C. Mean and SD are given, n>4 independent cultures for *E. coli* and n=6 for *Pseudomonas* species. The pMQ578 plasmid has *P*_*xut*_-*gfp*, and pMQ630 has *P*_*BAD*_-*gfp*. Xylose was used as an inducer for pMQ578 and arabinose was used for pMQ630.

Fold induction of GFP fluorescence was calculated for cultures with 10 mM compared to no inducer. For *P*_*BAD*_, the highest induction was for *E. coli* with a 46±36-fold increase compared to 19±3-fold and 27±9-fold for Pf0-1 and PA14, respectively. With *P*_*xut*_, GFP levels were unchanged for *E. coli* with a ratio of 1.03±0.2 for 10 mM compared to 0 mM xylose, but were 51±22-fold and 38±20-fold increased for Pf0-1 and PA14.

Utilization of an inducer as a carbon source by bacteria can reduce the efficacy of the inducer in a culture over time. We tested the ability of PA14 and Pf0-1 to grow in minimal medium with L-arabinose and D-xylose as a sole carbon source at 10 mM. Glucose at 10 mM was used as a positive control (Figure 4). After 24 hours of growth at 30°C, culture density was measured with a spectrophotometer. Pf0-1 grew with arabinose and glucose, but not xylose as a sole carbon source. PA14 grew only with glucose as a carbon source.

**Figure 4.**
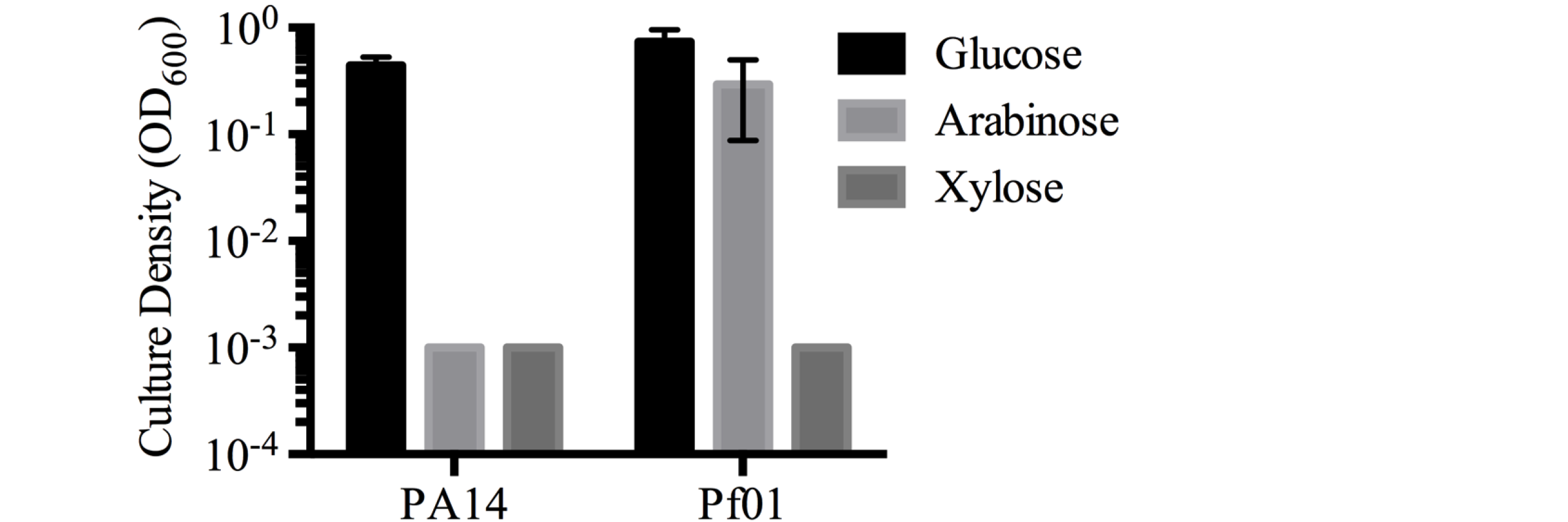
Tested Pseudomonas species do not use D-xylose as a sole carbon source. Cultures were incubated in M9 minimal medium with indicated sugars at 10 mM for 25 hours at 30°C. Mean and SD are given, n=6 independent cultures.

Ribose was also shown to be an inducer of *P*_*xut*_ in strain SBW25 (19); therefore, we evaluated ribose as an inducer (Figure 5). After 25 hours of growth at 30°C, PA14 showed a 19% increase in GFP fluorescence was observed with 10 mM ribose relative to no ribose, and a 93% increase was observed with 100 mM ribose (RFU: 1723±219 with 0 mM, 2050±265 with 10 mM, and 3335±464 with 100 mM). For Pf0-1, we observed a 25% increase of fluoresecence at 10 mM and a 41% increase at 100 mM ribose (RFU: 2074±172 with 0 mM, 2597±200 with 10 mM, and 2919±219 with 100 mM). This suggests that ribose would have limited utility as an inducer of *P*_*xut*_ relative to xylose in the two tested strains unless controlled, low levels, of gene expression are required for an experiment (Figure 5).

**Figure 5.**
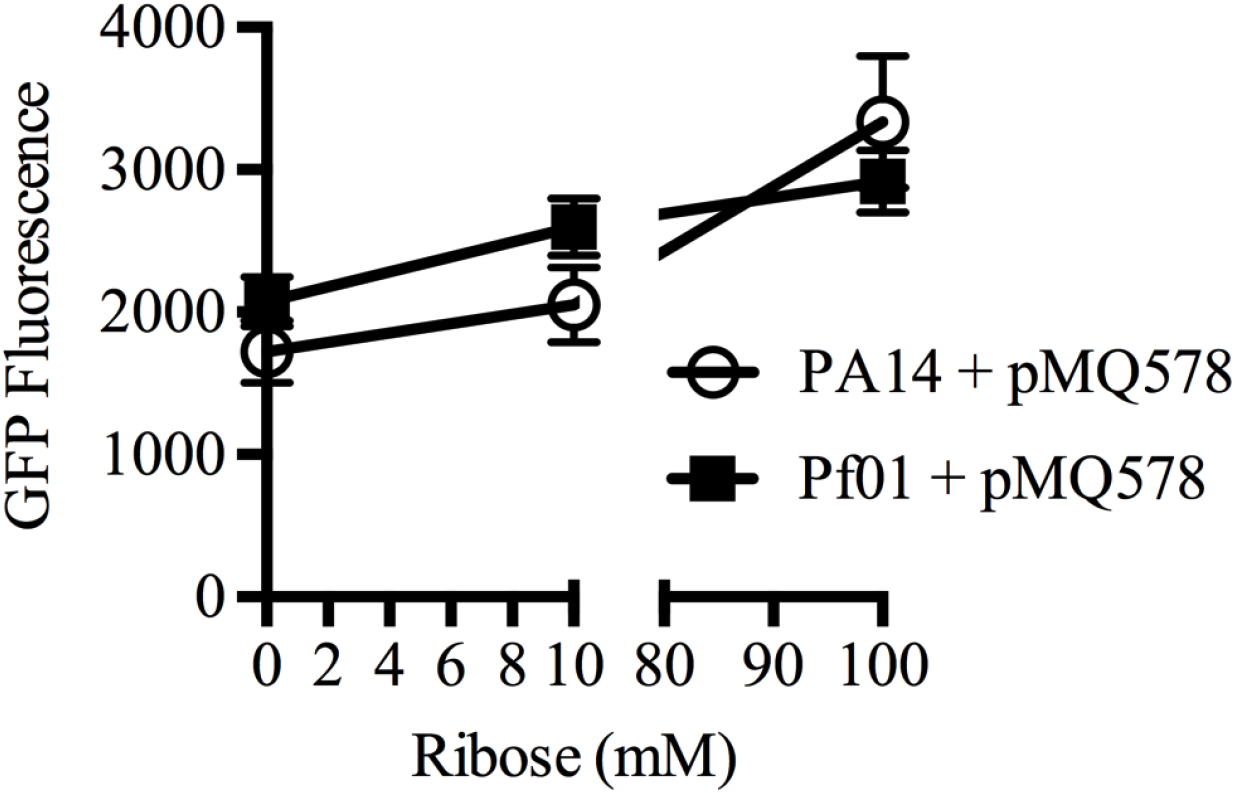
Evaluation of ribose for induction of *P*_*xut*_ promoter in two *Pseudomonas* species. Expression of *gfpmut3* from the two different promoters in a pBBRl-based plasmid. Cultures were prepared in LB medium with a range of inducer concentrations and grown at 30°C for 25 hours. Mean and SD are given, n=6 independent cultures. *P*_*xut*_ induction was minimally induced by ribose compared to xylose.

### Comparison of *P*_*BAD*_ and *P*_*xut*_ for induction of genes during stationary phase

We investigated whether these promoters were responsive to inducer for cultures in stationary phase (Figure 6). We tested the expression of *P*_*BAD*_-*gfp* and *P_xut_-gfp* in stationary phase cultures, to which inducer was added (10 mM) as a function of time. Pf0-1 and PA14 cultures were grown overnight, washed with PBS, and diluted to OD_600_=2 in PBS with inducer. A modest induction was shown by 6 hours in stationary phase cultures incubated in PBS compared to actively growing cultures (Figure 6). At the final time point, *P*_*xut*_ in Pf0-1 demonstrated the greatest induction of GFP fluorescence, with 5.5 ± 1.4 fold higher fluorescence than Pf0-1 with *P*_*BAD*_ (Figure 6). This suggests that the *P*_*xut*_ promoter could be used in stationary phase cultures, and perhaps biofilms, for expression of desired genes.

**Figure 6.**
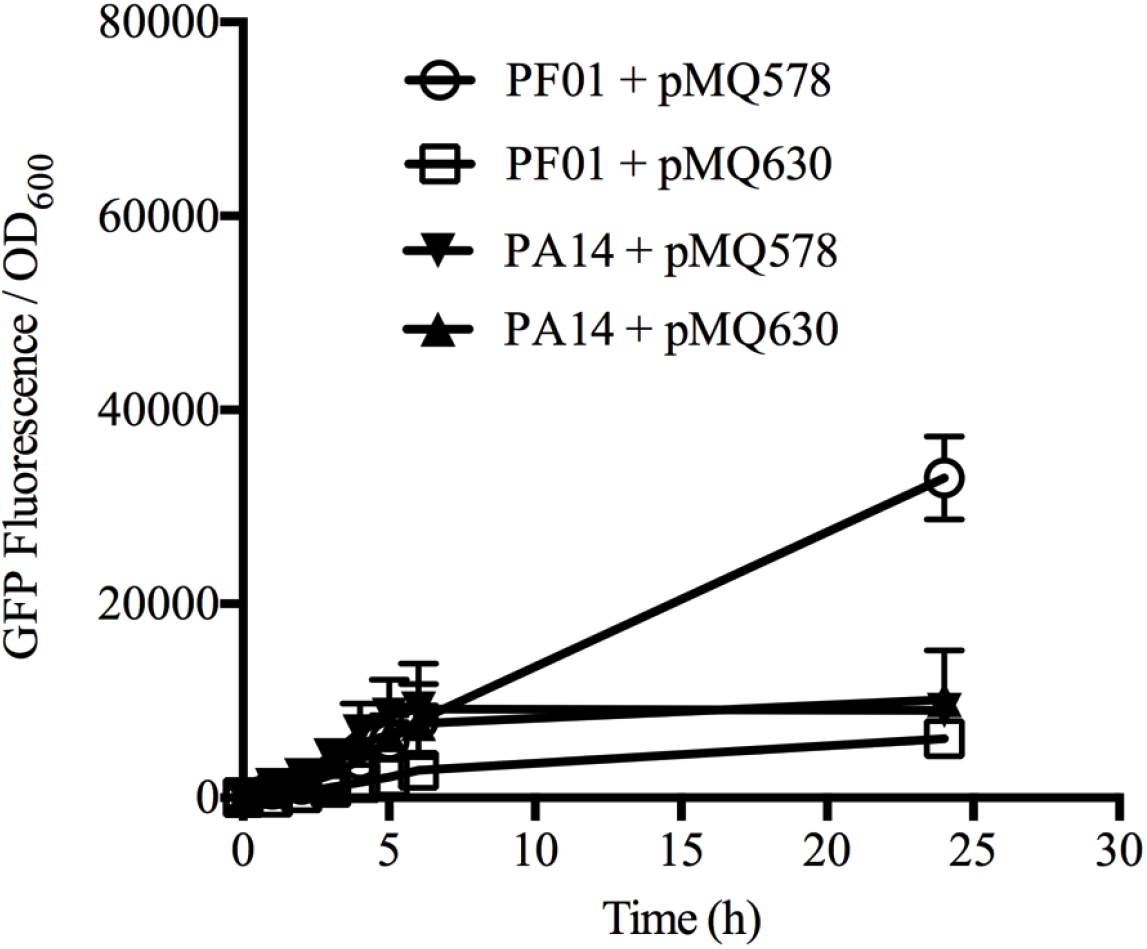
Stationary phase induction of *P*_*BAD*_ and *P*_*xut*_ promoter in *Pseudomonas* species. Bacteria were grown overnight in LB broth, washed, and adjusted to OD_600_=2.0 in PBS with inducer atlOmM. GFP fluorescence was measured over time. Mean and SD are given, n=3-4 independent cultures. *P*_*xut*_ was more highly inducible in *P. fluorescens* than in *P. aeruginosa* during stationary phase. *P*_*xut*_ and *P*_*BAD*_ were indistinguishable in *P. aeruginosa.* The pMQ578 plasmid has *P*_*xut*_-*gfp*, and pMQ630 has *P*_*BAD*_-*gfp*.

### Flow cytometry analysis of promoter activity in *Pseudomonas* species

Previous studies have indicated that the *P*_*BAD*_ promoter under many conditions is expressed in only a subset of cells of an induced population of *E. coli* (20–22). Here, we evaluated *P*_*BAD*_ and *P*_*xut*_ promoters in PA14 and Pf0-1 using flow cytometry analysis to assess promoter leakiness and to determine the frequency of cells which have gene protein production activated upon addition of a moderate level of inducer (10 mM). Data in Figure 7 and Figure 8A demonstrate a higher level of leakiness for *P*_*xut*_ compared to *P*_*BAD*_ in both PA14 and Pf0-1. For both species, the *P*_*xut*_ promoter produced higher fluorescence intensity in GFP-positive cells that were induced compared to *P*_*BAD*_, with a 2.5±8 and 2.3±0.5 fold for *P. fluorescens* and *P. aeruginosa* respectively (Figure 8B). A higher fluorescence intensity was observed for Pf0-1 with both promoters, but this may be due to the larger cell size of Pf0-1 compared to PA14 (p<0.001, ANOVA Tukey’s post-test). Cell length for PA14 was measured at 1.8 ± 0.5 or 1.6 ± 0.4 μm when grown in LB with xylose or arabinose at 10 mM respectively (n≥50 cells per group). Cell length for Pf0-1 was measured at 2.5 ± 0.7 or 2.4 ± 0.6 μm when grown in LB with xylose or arabinose at 10 mM respectively (n≥50 cells per group).

**Figure 7.**
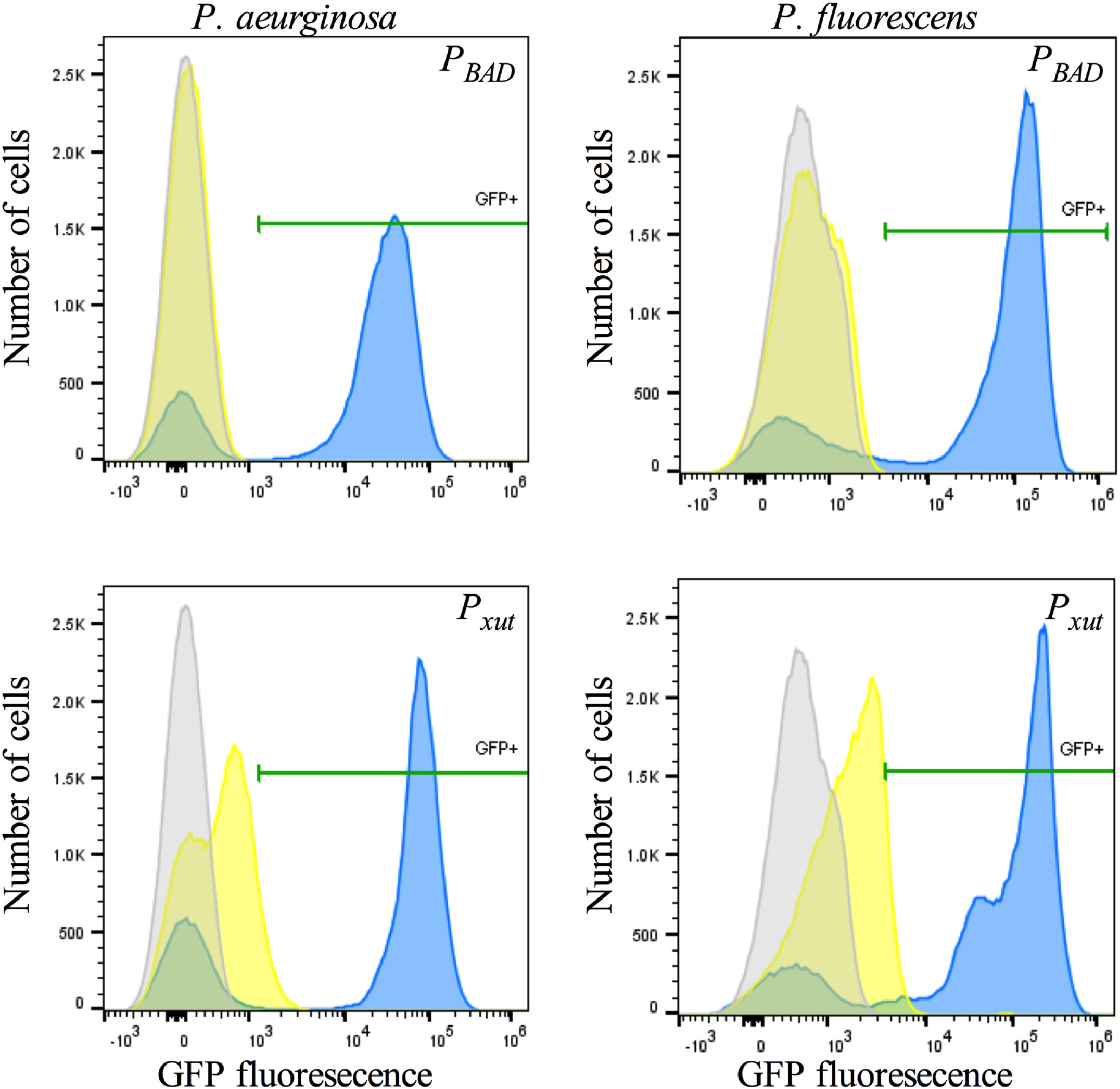
Flow cytometry analysis of *P*_*BAD*_ and *P*_*xut*_ promoter driven GFP expression in *Pseudomonas* species indicate increased leakiness and proportion of bacteria expressing GFP with inducer. Expression of *gfpmut3* from the two different promoters as measured by flow cytometry. Cultures were prepared in LB medium with inducer (10 mM) and grown at 30°C. The gray peak indicates PA 14 or PfOl with no plasmid to determine background levels. Yellow peaks are bacteria with pMQ578 or pMQ630 without inducer to indicate leakiness, and blue (and green) peaks are bacteria with pMQ578 and pMQ630 with inducer. The pMQ578 plasmid has *P*_*xut*_-*gfp*, and pMQ630 has *P*_*BAD*_-*gfp*. Xylose was used as an inducer for pMQ578 and arabinose was used for pMQ630. GFP fluorescence above the bacteria without plasmids were used to determine the positive cut-off noted as GFP+. A represenative experiment is shown (n = ~100,000 cells per group).

**Figure 8.**
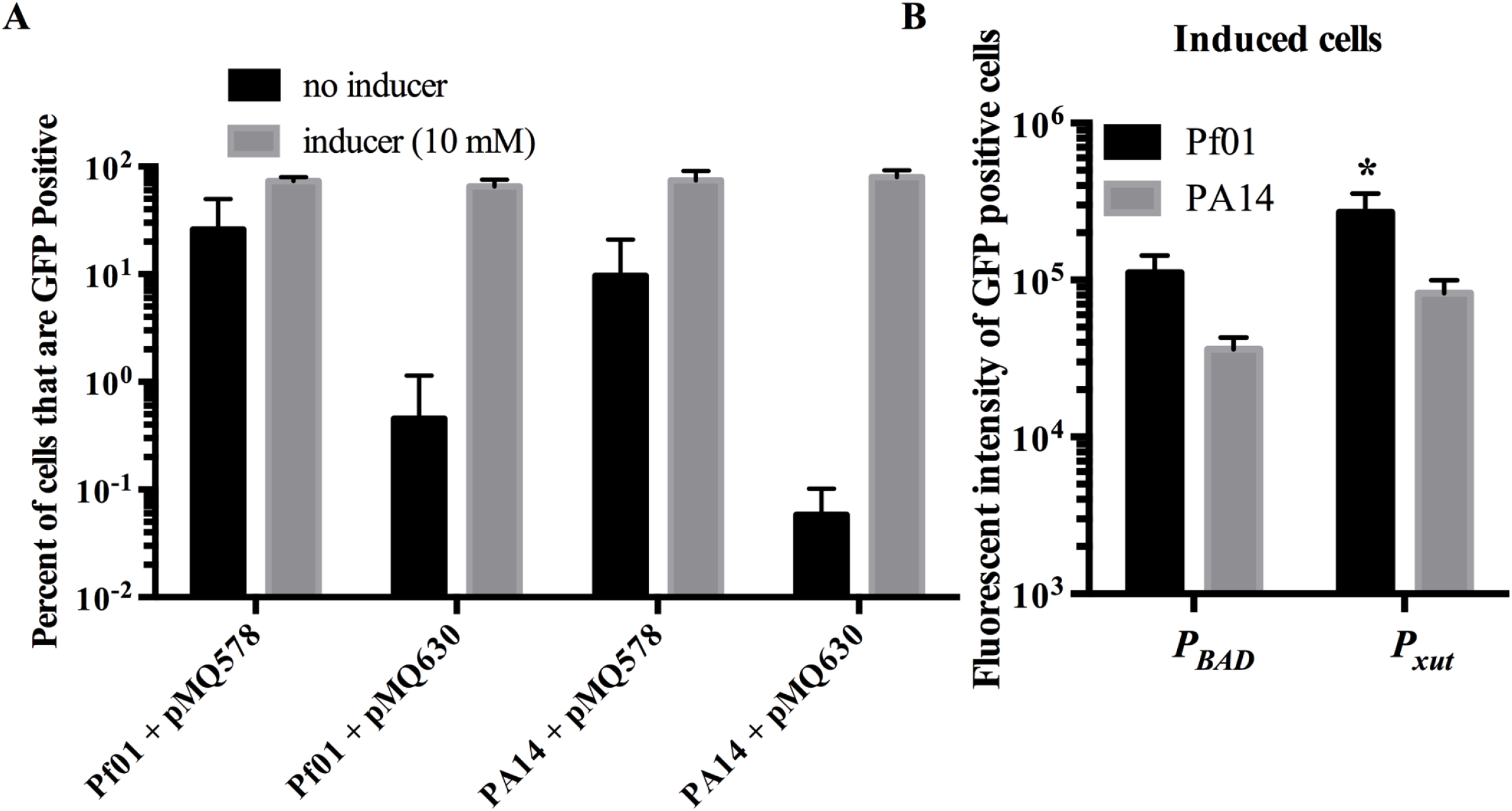
Analysis of promoter leakiness and strength using flow cytometry. Flow cytometry analysis (n=7) of GFP expression in *Pseudomonas* species. **A.** Percentage of GFP positive cells was determined relative to bacteria without a GFP plasmid. Mean and standard deviations are shown. pMQ578 has *P*_*xut*_ and pMQ630 has *P*_*BAD*_ drivign *gfp* expression. **B.** Fluorescent intensity of GFP positive cells that have been induced (10 mM inducer concentration). Means and standard deviations are shown. The asterisk indicates a significant difference from all other groups p<0.001 by ANOVA and Tukey’s post-test.

### Additional shuttle vectors with *P*_*xut*_

The pMQ578 vector with *P*_*xut*_-*gfp* did not have a convenient restriction site at the 5’ end of the *gfp* gene, which would impede replacing *gfp* with another gene using traditional cloning methods. To make more user-friendly constructs, the *gfp* gene was replaced with *tdtomato* flanked by restriction sites to make pMQ643 and with a multicloning site containing six unique restriction sites to make pMQ650 (Table 1). The pMQ643 plasmid was introduced into Pf0-1 and grown with and without xylose (10 mM). The culture with xylose was significantly brighter than the culture without xylose (p=0.016, unpaired T-test), with tdtomato fluorescence / OD_600_ for cultures with xylose measured at 1.3 × 10^5^ ± 7.2 × 10^4^ versus 2.0 × 10^3^ ± 1.7 × 10^3^ for the same culture conditions without xylose.

The pMQ650 plasmid was additionally modified by removing the yeast replicon and selectable marker for researchers who do not use yeast recombineering. The resulting plasmid, pMQ652, is 5.4 kbp rather than 7.3 kbp (Table 1).

### Use of the *P*_*xut*_ promoter system to evaluate the impact of *P. aeruginosa* corneal epithelial cell wound closure in vitro

A previous study demonstrated that secreted factors from *P. aeruginosa* strain PA14 could inhibit corneal cell migration and wound healing (12). To gain insight into the mechanism by which secreted factor(s) from *P. aeruginosa* inhibit corneal wound healing, we first heat-treated normalized secretomes from strain PA14. Whereas unheated secretomes inhibited wound healing, those heat treated for 10 minutes at 95°C were unable to inhibit corneal cell migration (data not shown). This result suggested that the inhibitory secreted factor was a heat labile molecule such as a protein.

*P. aeruginosa* has numerous secretion systems. We reasoned that the secreted inhibitor factor is unlikely to be secreted by type III or type VI secretion systems because contact between *P. aeruginosa* and the corneal cells was not necessary for the cell migration phenotype (only culture filtrates were used). Because many enzymes are secreted through the type II secretion system of *P. aeruginosa* (23), we tested whether the type II secretion system was required for inhibiting cell migration using a strain deficient in XcpQ. The XcpQ protein is an essential component of the type II secretion system and forms part of the outer membrane pore (23).

Unlike wild-type PA14 culture filtrates, those from a previously described Δ*xcpQ* derivative of PA14 (24) were unable to block cell migration (Figure 9). Expression of *xcpQ*, but not *gfp* (used as a negative control) from *P*_*xut*_, was able to restore the cell migration inhibition phenotype to the *xcpQ* mutant in a D-xylose-dependent manner; that is, 50, but not 5 mM D-xylose was sufficient to complement the cell migration inhibition phenotype (Figure 9). Importantly, as a control, D-xylose (50 mM) did not restore the cell migration inhibition phenotype to the Δ*xcpQ* mutant that did not have a plasmid indicating D-xylose alone was not responsible for the phenotype in the absence of the *xcpQ* plasmid (data not shown). These results indicate that the *P. aeruginosa* type II secretion system is necessary to inhibit corneal wound healing and demonstrates the utility of the *P*_*xut*_ system for studying *Pseudomonas* biology.

**Figure 9.**
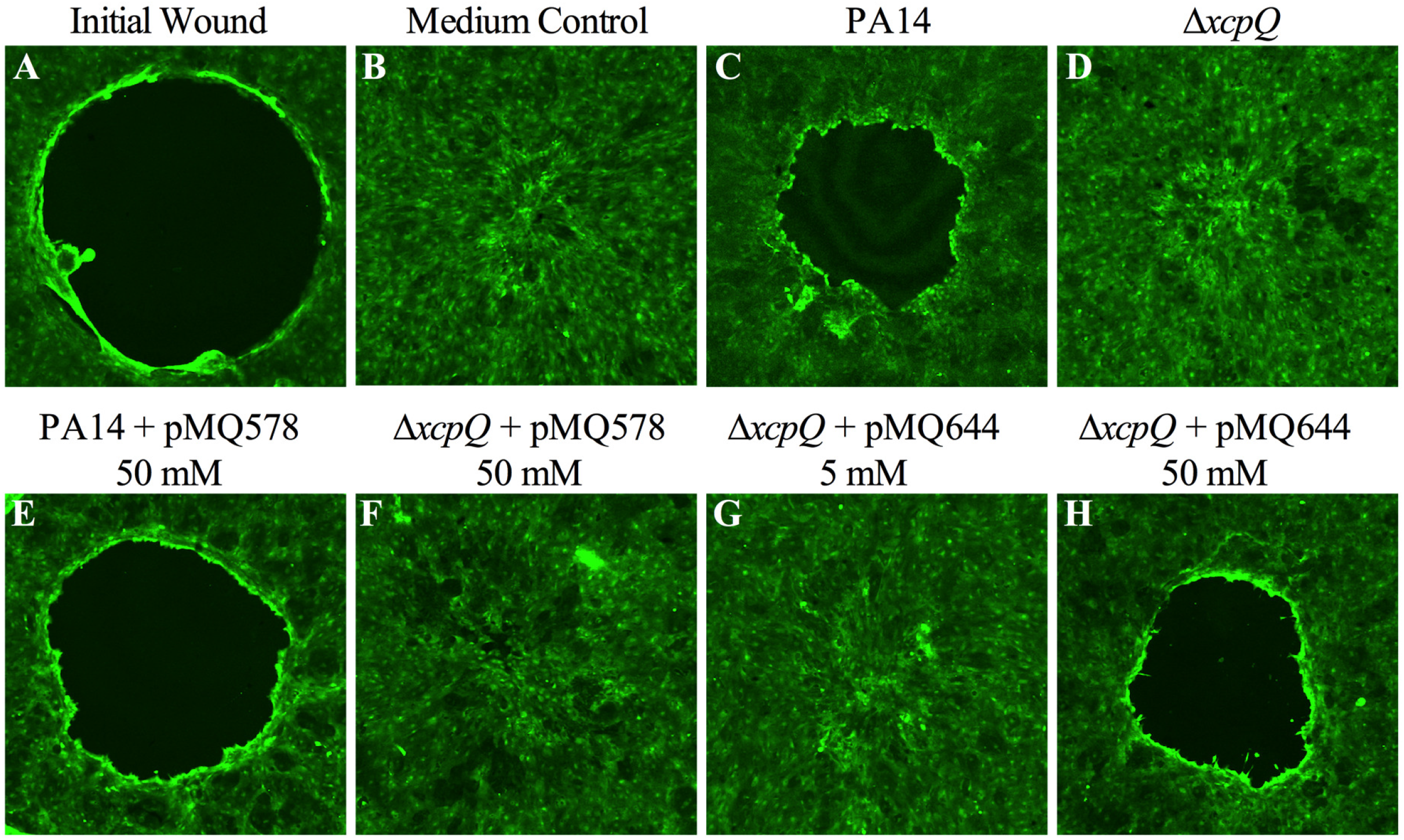
XcpQ is required for inhibition of stratified corneal cell migration in vitro. Reprentative images were stained with vital stain calcein AM and imaged by confocal microscopy. A cell free zone (black circle) in a layers stratified human comeal cell line **(A)** rapidly closes when treated with LB medium and incubated for 24 h **(B)**, but this migration is inhibited by culture filtrated from PA14 cultures grown in LB medium **(C)**. Filtrates from isogenic Δ*xcpQ* mutants were unable to impede cell migration **(D)**. D-xylose (50 mM) induced expression of gfp from pMQ578 did not alter the above noted migration phenotypes of cells exposed to PAM or Δ*xcpQ* filtrates **(E, F)**. Culture filtrates from the Δ*xcpQ* mutant with the wild-type *xcpQ* gene on a plasmid (pMQ644) that had been grown in LB medium with 5 mM D-xylose were unable to inhibit cell migration **(G)**, but those grown with 50 mM D-xylose were complemented for the cell migration inhibition defective phenotype **(H)**. All cultures were grown to stationary phase, adjusted to OD_600_=2 with fresh LB and bacteria were removed by centrifugation and filtration.

## DISCUSSION

This study introduces a new inducible promoter system for Pseudomonads. The new *P*_*xut*_ promoter had no detectable function in *E. coli*. This could be considered a negative of the *P*_*xut*_ promoter because one cannot use the same plasmid construct to express genes in *E. coli* and a *Pseudomonas* species. However, since plasmids are almost exclusively passed through *E. coli* before moving into *Pseudomonas* species, cloning of genes that are toxic to *E. coli* are likely to be more easily cloned with *P*_*xut*_ than other promoter systems that might have leaky expression in *E. coli*. For *P. aeruginosa*, the *P*_*xut*_ promoter appeared to be largely equivalent to *P*_*BAD*_ although it was leakier and tended toward higher expression with *P*_*xut*_ on a population basis (Figure 2) and in GFP-positive fluorescent cells, but did not reach significance (Figure 7 and 8). In *P. fluorescens*, the *P*_*xut*_ promoter performed better than *P*_*BAD*_ with respect to maximal expression levels and required a lower concentration of inducer for expression (Figure 2C, Figure 7, Figure 8). In addition to xylose, *P*_*xut*_ can be activated by xylulose and ribose (19) raising the possibility that the *P*_*xut*_ promoter can be fine-tuned with these alternative carbohydrate inducers.

*Pseudomonas* species have been suggested for industrial production and bioremediation of molecules including, but not limited to, β-peptides (25), polyhdroxyalkanoates (26), phenazine-1-carboxyamide (27), proteases (28, 29), pseudofactin (30), rhamnolipids (31–33), silver nanoparticles (34), and toluene (35). The ability of *P*_*xut*_ to activate expression of gene expression in the majority of cells and its relative strength in *P. fluorescens* compared to P_*BAD*_ suggest that it may be useful for larger scale gene synthetic applications. This is further strengthened by the low cost of D-xylose compared to other inducers, with L-arabinose and IPTG costing approximately 10-times more to induce a culture with 10 mM of the sugars or IPTG at 0.1 mM. An additional benefit of D-xylose is that it is not usable as a carbon source by either tested *Pseudomonas* species, indicating that the D-xylose inducer will not be catabolized for energy, and thereby eliminated from the culture over time.

With regard to corneal epithelial wound closure, previous work has implicated lipopolysaccharide from *E. coli* and *Serratia marcescens* in inhibiting corneal wound healing (12, 36). Unlike *P. aeruginosa*, the secreted inhibitory factor from *S. marcescens* was heat-stabile (12). Work with the *P*_*xut*_ promoter in this study strongly implicated the type II secretion system of *P. aeruginosa* strain PA14 in allowing the bacterium to inhibit corneal epithelial wound closure which may increase its ability to establish ocular infections.

## Acknowledgements

The authors would like to thank Nancy Baker for expert help with flow cytometry and Dr. George O’Toole at Dartmouth Medical School for gifts of strains. This study was supported by the Charles T. Campbell Laboratory of Ophthalmic Microbiology, NIH grant EY027331 (to R.M.Q.S.), and EY08098 (Core Grant for Vision Research), the Eye and Ear Foundation of Pittsburgh, and unrestricted funds from Research to Prevent Blindness.

